# Senolytic therapy combining Dasatinib and Quercetin restores the chondrogenic phenotype of human osteoarthritic chondrocytes by promoting the release of pro-anabolic mediators

**DOI:** 10.1101/2024.04.04.588076

**Authors:** Svenja Maurer, Valeria Kirsch, Leonie Ruths, Rolf E. Brenner, Jana Riegger

## Abstract

Cellular senescence is associated with various age-related disorders and is assumed to play a major role in the pathogenesis of osteoarthritis (OA). Based on this, we tested a senolytic combination therapy using Dasatinib (D) and Quercetin (Q) on aged isolated human articular chondrocytes (hACs), as well as in OA-affected cartilage tissue (OARSI grade 1-2). Stimulation with D+Q selectively eliminated senescent cells in both, cartilage explants and isolated hAC. Furthermore, the therapy significantly promoted chondroanabolism, as demonstrated by increased gene expression levels of COL2A1, ACAN, and SOX9, as well as elevated collagen type II and glycosaminoglycan biosynthesis. Additionally, D+Q treatment significantly reduced the release of SASP factors (IL6, CXCL1). RNA sequencing analysis revealed an upregulation of the anabolic factors, inter alia, FGF18, IGF1, and TGFB2, as well as inhibitory effects on cytokines and the YAP-1 signaling pathway, explaining the underlying mechanism of the chondroanabolic promotion upon senolytic treatment. Accordingly, stimulation of untreated hAC with conditioned medium of D+Q-treated cells similarly induced the expression of chondrogenic markers. Detailed analyses demonstrated that chondroanabolic effects could be mainly attributed to Dasatinib, while monotherapeutical application of Quercetin or Navitoclax did not promote the chondroanabolism. Overall, D+Q therapy restored the chondrogenic phenotype in OA hAC most likely by creating a pro-chondroanabolic environment through the reduction of SASP factors and upregulation of growth factors. This senolytic approach could therefore be a promising candidate for further testing as a disease-modifying osteoarthritis drug.

## Introduction

Osteoarthritis (OA) is the most prevalent joint disease worldwide. It presents a complex disorder of the entire joint, including cartilage destruction, sclerosis of subchondral bone, formation of osteophytes and synovitis [1–3]. The clinical pattern is characterized by pain and joint stiffness [4]. So far, OA is considered as an uncurable disease and the therapy is mainly limited to symptom relief [5]. In recent years, much research has been conducted in developing disease modifying OA drugs (DMOADs), comprising mainly anti-inflammatory, anti-catabolic and chondroanabolic strategies. Recently, cellular senescence has emerged as a potentially promising target in OA therapy [6–8]. Cellular senescence is characterized by an irreversible cell cycle arrest and was first discovered in fibroblasts by Hayflick and Moorhead in 1961 [9]. Besides its relevance as a potent anticancer mechanism and its importance during embryogenesis and wound healing, cellular senescence is one of the hallmarks of aging and associated with several age-related diseases [1, 10]. In line with this, an accumulation of senescent cells was discovered in OA tissue, which might be linked to the pathogenesis of the disease. It is thought that stress factors, such as oxidative stress, release of damage associated molecular patterns (DAMPs) or chronic inflammation after joint trauma or during aging play an important role in the accumulation of senescent cells in OA, consequently referred to as stress-induced premature senescence (SIPS) [1, 11, 12]. Senescent chondrocytes display an upregulation of anti-apoptotic pathways by an increase of B-cell lymphoma 2 and -XL family (BCL-2, -XL), enhanced expression of the cyclin dependent kinase inhibitors p21^Cip1/WAF1^ (CDKN1A) and p16^INK4a^ (CDKN2A), elevated activity of the senescence associated β-galactosidase (SA-β-Gal), and are characterized by a so-called senescence-associated secretory phenotype (SASP) [12–16]. Senescent chondrocytes secrete elevated levels of pro-inflammatory chemokines and cytokines, such as CXCL-1, IL-1, IL-6 and IL-8, but also matrix metalloproteinases like MMP-1, -3-, and -13. These SASP factors contribute to cartilage degeneration and “inflammaging”, which can be understood as the spreading of senescene to neighboring cells due to chronic inflammation [1, 12, 13, 16]. Overall, senescent chondrocytes represent a potential therapeutic target. Consequently, senotherapeutics play an increasingly important role in the research field of OA. The senolytic drugs are divided into senomorphics and senolytics. While senomorphics target the secretion and action of SASP factors, senolytics eliminate senescent cells by the selective induction of apoptosis [16, 17].

Among the senolytic drugs, the combination of Dasatinib and Quercetin (D+Q) has turned out to be particularly efficient in the context of musculoskeletal diseases as proven in murine [18] and rat models [19] of OA, bone healing [20–22], and disc degeneration [23]. Dasatinib is an already FDA and EMA approved drug (Sprycel®) for the treatment of chronic myeloid leukemia (CML) and philadelphia-chromosome positive acute lymphoblastic leukemia (Ph+ALL) [24]. The drug exerts its pro-apoptotic effects through inhibition of several tyrosine kinases [24, 25], which contribute to the pro-survival network in cancer and senescent cells [23, 26, 27]. Quercetin on the contrary is a flavonoid, found in fruits and vegetables, such as grapes, citrus fruits, onions, etc. and is known for its anti-inflammatory, antioxidative, anti-cancer, and anti-microbial effects [28–30]. It exerts its pro-apoptotic effects in senescent cells by inhibition of the phosphatidylinositide-3-kinase (PI3K)/Akt pathway and downregulation of BCL-2 [29, 31]. Although D+Q efficiently attenuated progression of OA after distraction arthroplasty in rats [19] and reduced age-related temporomandibular joint (TMJ) degeneration in mice [18], none of the previous studies investigated the effects of D+Q on human OA chondrocytes. Moreover, the phenotype of surviving chondrocytes after D+Q treatment has not been addressed so far. To close this gap in research, we investigated the efficacy of the combined application of the senolytic agents D+Q on OA hAC and cartilage tissue to particularly evaluate the chondrogenic characteristics after elimination of senescent cells. Further, we compared the outcome of D+Q treatment with one of the most commonly used senolytic, Navitoclax, which exerts its senolytic properties through the inhibition of BCL-2/-XL [32–34].

Our results indicate that the treatment with D+Q selectively eliminates senescent cells in human OA cartilage tissue and in isolated hAC. The therapy decreased the expression of SASP factors and enhanced that of growth factors IGF1, FGF18, and TGFB2. As a result, D+Q restored the chondrogenic phenotype of OA hAC as demonstrated by a remarkable increase of collagen type II (COL2) and proteoglycan (PG) production. Accordingly, the therapy also resulted in a recovery of the chondrogenic redifferentiation potential of high passage chondrocytes. While the chondroanabolic effects could be mainly attributed to Dasatinib, the senolytic Navitoclax impaired the chondrogenic phenotype of hAC. Moreover, selective elimination of senescent cells and increase of COL2 synthesis was confirmed in OA cartilage tissue. Overall, this is the first study demonstrating that D+Q therapy, and in particularly Dasatinib, stabilizes and possibly even restores the chondrogenic phenotype of human OA chondrocytes. These novel findings provide important insight about the divergent therapeutic efficacy of senolytics in OA and are of particular interest regarding future application of D+Q in clinical studies.

## Methods

### Cultivation of Human Cartilage Tissue and Isolation of Chondrocytes

Human articular cartilage tissue was obtained from donors undergoing knee replacement surgery. Informed consent was received from patients according to the regulations of the Ethics Committee of the University of Ulm (ethical approval number 353/18). Cartilage explants were created using a biopsy punch (⌀ 6 mm) from macroscopically intact tissue (OARSI score ≤ 2) of human femoral condyles and thereafter cultured in serum-free medium in an incubator (37°C, 5% CO_2_ and 95% humidity) (S1a) [35]. Isolation of chondrocytes was performed by cutting cartilage slices from condyles, digesting with 0.2% protease (Sigma-Aldrich) for 45 min at 37°C and further digesting overnight in 0.025% collagenase (collagenase from clostridium histolyticum, Sigma-Aldrich) at 37°C. Afterwards, chondrocytes were filtered (cell strainer 40 µm), centrifuged, resuspended and cultured in serum-containing medium (S1a). Passaging was performed at approximately 80% confluence [12]. Cells were used at low (P≤2) and high passages (P≥4) as indicated in the figure legends.

### Porcine Cartilage Tissue

Porcine knee joints were obtained from a butcher (age: 4-6 months). Cartilage tissue was harvested from femoral condyles and cultured in serum-free medium containing high glucose DMEM (S1a).

### Senolytic treatment

During running experiments, isolated chondrocytes were cultured in serum reduced medium (5% FBS), consisting of 1:1 serum-containing medium and serum-free medium (S1a). Cells were treated either for 3 d with Dasatinib (50-500 nM) (S1021, Seleckchem) and Quercetin (5 µM) (S2391, Seleckchem) or with Navitoclax (5 µM) (S1001, Seleckchem) followed by 4 d without senolytics (d7). Untreated cells served as control. Senolytics were dissolved in DMSO, hence, a DMSO control was included. Medium was changed at least three times a week. Supernatants were collected at d3 and 7 and total RNA was isolated. In contrast to isolated chondrocytes, cartilage explants were cultured in serum-free medium (S1a) during the experiments and stimulated according to the same application scheme, including additional analysis at d14.

### Stimulation with SIPS- and SASP-Conditioned Medium

hAC were treated with 0.1 µM Doxorubicin (Doxo) (Selleckchem) for 7 d to create a stress-induced premature senescence (SIPS) [12]. Next, the supernatant of untreated and treated (SIPS-CM) cells was collected 48 h after deprivation of Doxorubicin. Supernatants were used to stimulate hAC, in order to study paracrine effects.

Furthermore, high passage hAC were treated for 3 d with D (250 nM) + Q (5 µM) or cultured without senolytic treatment. Afterwards, cells were cultured for 2 d without the senolytics and medium was collected at d5. Cells were split and cultured with medium from previously untreated cells (SASP-CM) or with medium from cells prior treated with D+Q (DQ-CM). Cells cultured with fresh medium (fresh-M) served as control. Overall, this experiment includes the following stimulation groups: group 1 (5 d w/o treatment, 3 d with fresh-M), group 2 (5 d w/o treatment, 3 d with SASP-CM), group 3 (5 d w/o treatment, 3 d with DQ-CM), group 4 (3 d with D+Q treatment, 2 d w/o treatment, 3 d with fresh-M), group 5 (3 d with D+Q treatment, 2 d w/o treatment, 3 d with SASP-CM) and group 6 (3 d with D+Q treatment, 2 d w/o treatment, 3 d with DQ-CM) (Fig. 6h).

### ATDC5 Murine Chondrogenic Progenitor Cell Line

ATDC5 cells were cultured in serum containing medium (S1a) [36]. During the experiments, cells were either cultured in chondrogenic differentiation medium (CDM) (S1a) for 7 d, or in serum-reduced medium with D (250 nM) + Q (5 µM) for 3 d. In a further experiment, the cells were cultured with D+Q stimulation and with or w/o ITS (3 d). All analyses were performed at d7 and the medium was changed at least three times a week.

### Cytotoxicity/Cell Proliferation Assay – alamarBlue^TM^

This cell viability assay is based on the ability of healthy cells to metabolize the blue dye resazurin into the red fluorescent resorufin. After treatment with the senolytics, cells were treated with 5% Resazurin solution for 3 h at 37°C. Afterwards, the fluorescence intensity (FI) was measured using a microplate reader (Tecan, Microplate Reader infinite M200 pro) with an excitation (ex) wavelength of 550 nm and emission (em) detection at 590 nm [12].

### Caspase3/7 Activity Assay

Caspase-3 activity as an important initiator of apoptosis was assessed by the Amplite^TM^ Fluorimetric Caspase 3/7 Assay Kit (AAT Bioquest) and performed according to the manufacturer’s instruction. Cells were stimulated for 6 h with Navi (2.5-5 µM) and treated for 2 h with the Caspase 3/7 working solution at room temperature (RT) in the dark. Afterwards, the plate was centrifuged and fluorescence was measured with a microplate reader (ex= 350 nm, em= 450 nm) [12].

### Cell Viability Assay – Live/Dead Staining

To determine cell viability in cartilage explants, the LIVE/DEAD™ viability/cytotoxicity kit for mammalian cells (Invitrogen) was used. Live cells were stained green with calcein AM (1 µM), while dead cells were stained red with ethidium homodimer-1 (2 µM). In short, staining of 0.5 mm thick cartilage slices for 30 min and microscopic image taking by means of a z-stack (Zeiss, Microscope Axioskop 2 mot plus, Software AxioVision) [35].

### Gene Expression Analysis – qPCR

Total RNA was isolated from hAC using the RNeasy Mini Kit (Qiagen) and afterwards reverse transcribed to cDNA by means of the Superscript II Reverse Transcriptase (Invitrogen). Gene expression levels were determined by quantitative real-time polymerase chain reaction (RT-PCR) (StepOnePlus^TM^ Real-Time PCR System; Applied Biosystems) with the 2^-ΔΔCT^ method [35]. GADPH, HPRT1 and in case of hAC additionally TMEM199 was used as reference genes. Target genes were analyzed by the TaqMan^TM^ Gene Expression Assay (S1b).

### RNA-Sequencing Analysis

Total RNA of hAC of three donors w/o or with 250 nM D + 5 µM Q therapy for 3 d was sent to Biomarker Technologies (BMK) GmBH (Münster, Germany) for mRNA sequencing analysis. Bioinformatic analysis and generation of graphics were performed by the company.

### Supernatant Analysis – ELISA

Supernatants of cultured hAC and cartilage tissue were harvested and analyzed by means of enzyme-linked immunosorbent assays (ELISAs). SASP factors: human IL-8 Uncoated ELISA (88-8086-88, Invitrogen), human IL-6 Uncoated ELISA (88-7066-88, Invitrogen) and human GRO alpha (CXCL1) Uncoated ELISA (88-52122-22, Invitrogen). Collagen type II synthesis: Procollagen II C-Propeptide (CPII) ELISA (60-1003-001, IBEX Pharmaceuticals Inc.). All steps were performed according to the manufacturers’ instruction [12, 37]. As treatment with senolytics eliminates senescent cells, raw ELISA data of the SASP factors were normalized to the alamarBlue values.

### Immunofluorescence Staining (IF): COL2 and Ki67

hAC were fixed with 4% formalin and blocked for 1 h at 37°C (Protein-Block serum free, X0909 Dako Agilent). COL2 antibody (MA5-13026, Invitrogen) was diluted 1:100 (Antibody Diluent, S0809 Dako Agilent) and cells were incubated for 2 h at RT. Afterwards, secondary biotinylated antibody (biotinylated link, K0690 Dako Agilent) and the corresponding dye iFluor^TM^568-streptavidin (AAT Bioquest) were used. Nuclei staining was performed using Dapi (0.25 µg/µL). IF staining was completed by mounting the cells (Fluorsave^TM^ Reagent, Merck) and capturing images under a fluorescent microscope (Zeiss, Microscope Axioskop 2 mot plus). FIs were analyzed using the software ImageJ2 (version 2.3.0/1.53q) with a modified CTCF method, taking into account the cell number (CTCF = (integrated density [image] – (area [image]*mean background fluorescence))/cell number). In case of Ki-67 staining, cells were after fixation permeabilized with 0.1% triton-x-100 for 15 min at RT, blocked for 1 h and the overnight staining was performed with the primary antibody (ab16667, dilution 1:250) at 4°C. Continuation with secondary antibody Alexa Fluor488 (ab15077) for 30 min at RT, and Dapi staining for 15 min at RT.

### Alcian Blue Staining

Glycosaminoglycans (GAGs) were stained by using alcian blue. hAC were fixed with 4% formalin followed by incubation with 0.1 M HCL (Merck) for 10 min. Subsequently, staining was performed with 1% alcian blue solution (Roth) for 30 min during shaking. Cells were washed three times with 0.1 M HCL and images were taken by means of a light microscope (Zeiss, Axiovert 35). In case of ATDC5 cells, destaining with 1% sodium dodecyl sulfate (SDS) (Sigma Aldrich) was performed and absorbance was measured at 595 nm.

### SA-β-Gal Activity Assay

For measuring the SA-β-Gal activity, the 96-Well Cellular Senescence Assay Kit (Cell Biolabs) was used. In short, 5000 cells were seeded onto the wells. After 24 h, cell lysis was performed using the cell lysis buffer. Subsequently, cell lysates were centrifuged and the supernatant was collected, and incubated with the assay buffer at 37°C for 2 h. Stopping solution was used for termination of the reaction and fluorescence intensity was measured (ex= 360 nm; em= 465 nm) [38].

### Chondrogenic Redifferentiation of hAC

High passage hAC were treated with the senolytics for 3 d and subsequently cultured in a 3D pellet culture (3.5×10^5^ cells) for 28 d in CDM. During the first 7 d, the growth factors rhBMP-6 and rhTGF-β3 were added freshly every day and afterwards with medium change thrice a week [39]. Chondrogenic differentiation potential was assessed by an adopted scoring system as previously described, including the pellet size, staining intensities of safranin-O and COL2 (IHC), cell morphology and the produced matrix (cell-cell distance)[40].

### Histology and Immunohistochemistry

Cartilage explants and chondrocyte pellets were fixed with 4% formalin, dehydrated, embedded in paraffine, and cut in 3.5 µm thick sections to perform IHC and safranin-O staining. Briefly, samples were dewaxed and rehydrated, Weigert’s iron hematoxylin was used for nuclear staining, safranin-O (Chroma-Gesellschaft) for staining of PGs and fast-green (Sigma Chemical Company) as a counterstaining. For the COL2-IHC staining, samples were as well dewaxed, rehydrated and then digested for 40 min at 37°C with pepsin (Sigma Aldrich). In case of Ki67 and CDKN2A, samples were incubated in citrate buffer overnight for antigen-retrieval. 3% H_2_0_2_ treatment was performed for blocking of endogenous peroxidase and further protein blocking with Dako-blocking buffer was performed. Next, samples were incubated overnight with the corresponding primary antibodies (COL2-AB 1:2000, DSHB II-II6B3-s; Ki-67-AB 1:200, ab16667; CDKN2A-AB 1:200, LS-B1347) at 4°C. Staining was continued using the Dako LSAB2 System-HRP kit (Dako Agilent) and Mayer’s hematoxylin (Merck) solution was used for nuclei staining [39].

### Statistical Analysis

Data sets were analyzed using GraphPad Prism 10 software. Outliers were quantified for data sets n≥5 by means of a Grubbs’ test and excluded from statistical analysis. The comparison of two groups with each other was performed using a t-test, whereas a multiple t-test, one-way or two-way ANOVA was performed for the comparison of several groups (significance level α= 0.05). Detailed statistical information can be obtained from the corresponding figure captions.

## Results

### D+Q Therapy Exhibits Cytotoxic Effect on High-Passage hAC

To investigate senolytic effects of D (25-500 nM) and Q (0-50 µM) on OA hAC, we first performed a broad screening of different concentrations of the drugs and determined cytotoxicity by means of an alamarBlue assay. The results demonstrate that higher concentrations of D and Q are associated with a reduction in the FI, which indicates cytotoxic effects of D+Q treatment (Fig. 1a). Based on these results, an optimal treatment range of 25-500 nM D in combination with 5 µM Q was identified and thus chosen for the following experiments.

**Figure 1:**
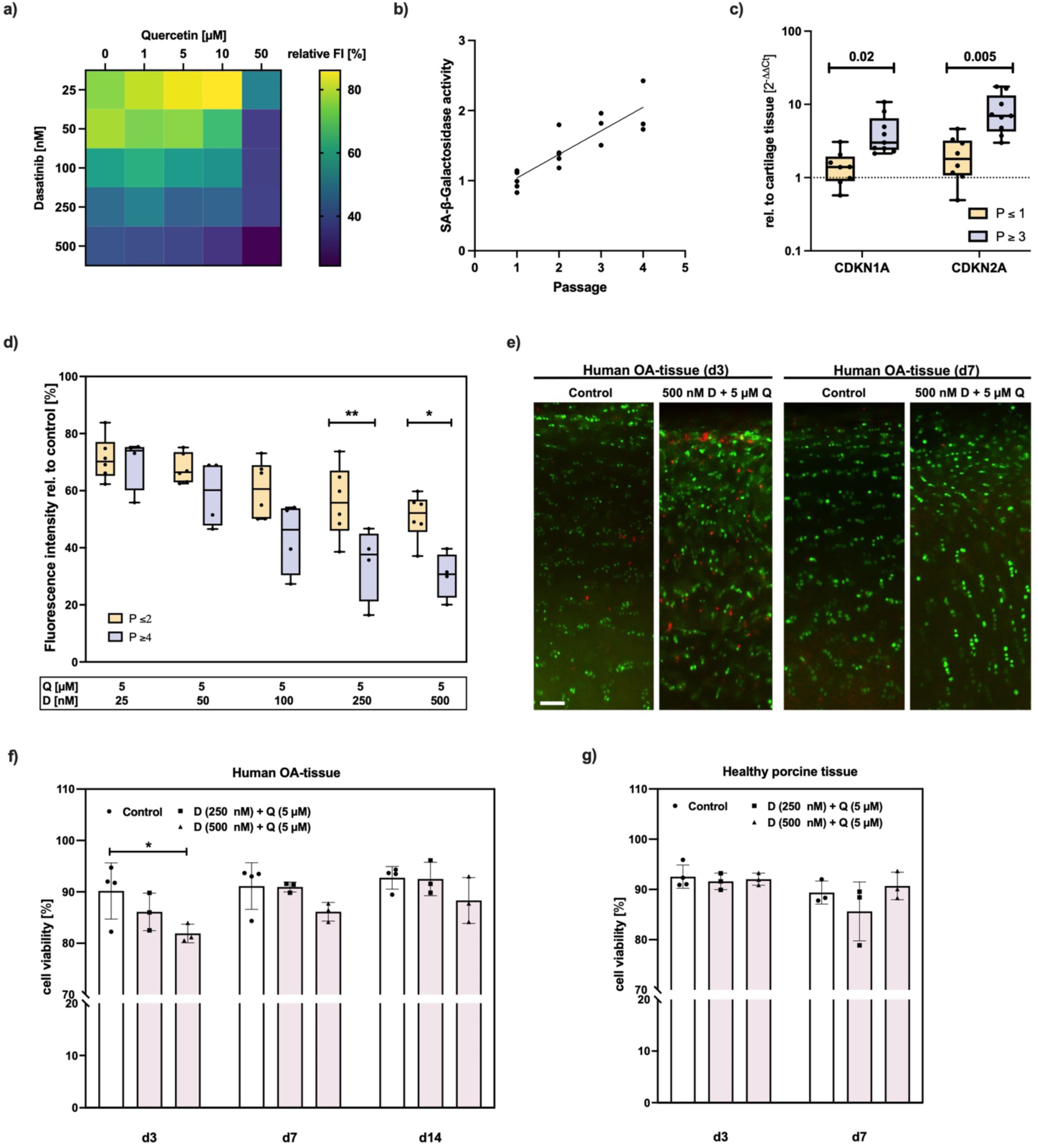
Selective elimination of senescent hAC by the senolytics D+Q. a) alamarBlue assay of isolated hAC after D (25-500 nM) + Q (0-50 µM) treatment for 3 d. b) SA-β-Gal activity of isolated hAC (passage 1-4) and c) mRNA levels of CDKN1A and -2A of hAC at low vs. high passage. d) alamarBlue assay of low vs. high passage hAC after D+Q stimulation for 3 d. e) Exemplary live/dead images of human cartilage tissue w/o and with D+Q stimulation for 3 d at d3 and d7 (scale bar = 100 µm). f) Corresponding cell viability analysis of human (OARSI grade 1-2) and g) healthy porcine cartilage after treatment for 3 d with D+Q (analysis at d3, d7 and for human cartilage additionally at d14). Statistical analysis: c) multiple unpaired t test; d) two-way ANOVA with *Šidák*’s multiple comparisons test: *p<0.05, **p<0.01; f) two-way ANOVA with *Dunett’s multiple* comparisons test: *p<0.05.

Next, we determined whether cell toxicity of D (25-500 nM) and Q (5 µM) on OA hAC resulted from selective senolysis of senescent cells. For this purpose, we first confirmed the increase of cellular senescence through passaging by means of an SA-β-Gal activity assay, which displayed a correlation between passaging and the enzymes’ activity (Fig. 1b). Moreover, passaging resulted in an altered cell morphology, demonstrated by an elongated cell shape (S2a-b). The accumulation of senescent cells was further confirmed on gene expression level. In the following, OA hAC of passage ≥4 are referred to as “high passage hAC”, while hAC of passage ≤2 as “low passage hAC”. High passage hAC expressed significantly elevated CDKN1A and CDKN2A mRNA levels as compared with low passage hAC (Fig. 1c). Stimulation of high and low passage hAC with D+Q resulted in a significantly more pronounced reduction of the FI in cells of high passages in case of 250 nM and 500 nM D (Fig. 1d). Therefore, these concentrations were used for all subsequent experiments. Moreover, this finding provides first evidence that D+Q might selectively eliminate senescent cells. To further investigate the selective removal of senescent cells, human OA cartilage exhibiting moderate degeneration (OARSI grade 1-2) and healthy porcine cartilage tissue (OARSI grade 0) was treated with the senolytics. By means of a live/dead staining, it was demonstrated that the senolytic treatment resulted in increased cell death in OA tissue (Fig. 1e+f), whereas no cytotoxic effects were observed in healthy porcine cartilage (Fig. 1g). Cell viability in human explants recovered after removal of the therapeutics (Fig. 1f).

### D+Q Treatment Reduces Cell Proliferation in Isolated hAC and Percentage of p16 Positive Cells in Cartilage Tissue

It is assumed that the cell cycle regulator p21^WAF1/Cip1^ (CDKN1A) is required for the initiation of cellular senescence and p16^INK4a^ (CDKN2A) for maintaining this state [41]. Therefore, their expression was studied in greater depth directly after senolytic treatment and 4 d after withdrawal of the therapeutics. Directly after therapy with D+Q (d3), no decrease of CDKN1A mRNA levels was found in high passage hAC (Fig. 2a), whereas the expression was significantly downregulated 4 d after withdrawal of the senolytics (d7) (Fig. 2b). CDKN2A mRNA levels were not affected by the senolytics *in vitro* (Fig. 2c+d). Treatment of isolated hAC with D+Q resulted in a reduction of Ki-67 positive cells 2 d after deprivation (Fig. 2e). These results are in line with the alamarBlue assay, in which D+Q treated cells exhibited a lower growth rate after withdrawal of the senolytics (Fig. 2f). In cartilage tissue, only few cells were positive for Ki-67 and no significant differences were found at d7 and d14 in D+Q-treated tissue as compared to the untreated control (Fig. 2g). In contrast to the *in vitro* analysis, D+Q stimulation resulted in a distinct reduction of CDKN2A positive cells in moderately degenerated cartilage tissue from OA patients (OARSI grade 1-2) (Fig. 2h+i), indicating the selective clearance of senescent cells.

**Figure 2:**
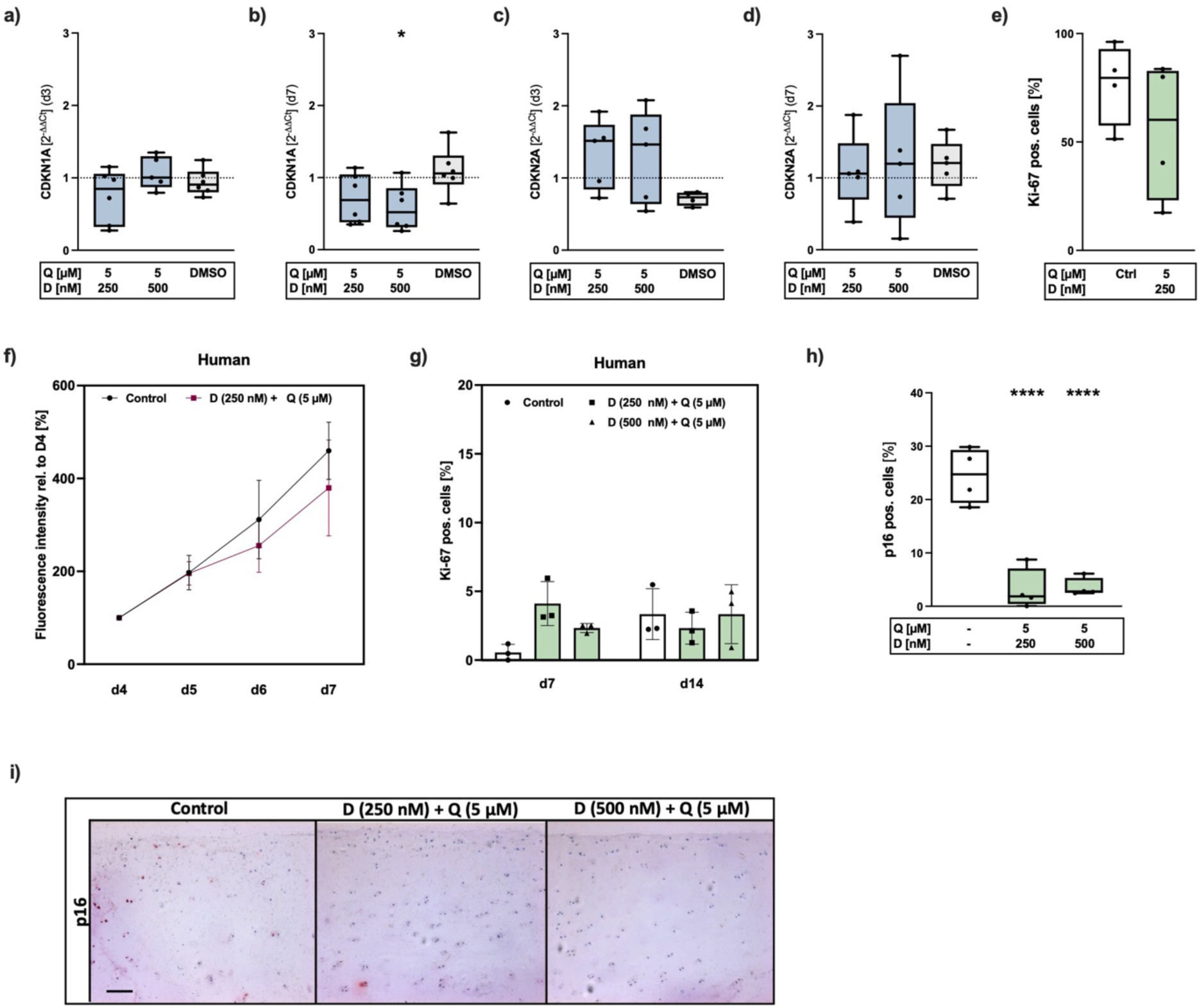
Influence of D+Q therapy on cell cycle regulators and cell proliferation. a)+b) mRNA levels of CDKN1A and c+d) CDKN2A of high passage hAC during (d3) and after deprivation (d7) of D+Q. e) Percentage of Ki-67 positive cells in isolated hAC (IF staining) and f) the measured cell proliferation by means of an alamarBlue assay relative to d4. g) Ki-67 positive cells in cartilage tissue, h) quantification of p16 positive cells and i) the exemplary images (IHC) after D+Q treatment of human cartilage explants (scale bar = 200 µm). Blue colored box plots = gene expression analysis data; green colored box plots = data of protein analysis. Statistical analysis: b) one-way ANOVA with *Šidák*’s multiple comparisons test: *p<0.05; h) one-way ANOVA with Dunnett’s multiple comparisons test: ****p<0.0001.

### D+Q Treatment Promotes Chondroanabolism

It is well known that chondrocytes rapidly lose their chondrogenic phenotype during *in vitro* culturing, including aggrecan and collagen type II expression among other characteristics [42, 43]. For this reason, we have investigated the effects of D+Q on the chondroanabolism of high passage hAC. The gene expression of ACAN was significantly upregulated directly after therapy and 4 d after withdrawal of the drugs (Fig. 3a+b). Interestingly, COL2A1 expression was likewise significantly increased at d3, but even further enhanced, from about 7-fold (d3) to approximately 30-fold, after deprivation of the senolytics (Fig. 3c+d). Moreover, D+Q treatment resulted in a significant increase of SOX9 mRNA levels directly after therapy and were still slightly enhanced after withdrawal (Fig. 3e+f). Asides from that, higher amounts of GAGs and a significantly increased biosynthesis of COL2 were detected in 2D-cultured isolated hAC (Fig. 3g-i). In line with the cell culture experiments, we could confirm elevated COL2 production in human cartilage tissue by means of a CPII ELISA (Fig. 3j+k) and IHC staining against COL2 (Fig.3l) (S3).

**Figure 3:**
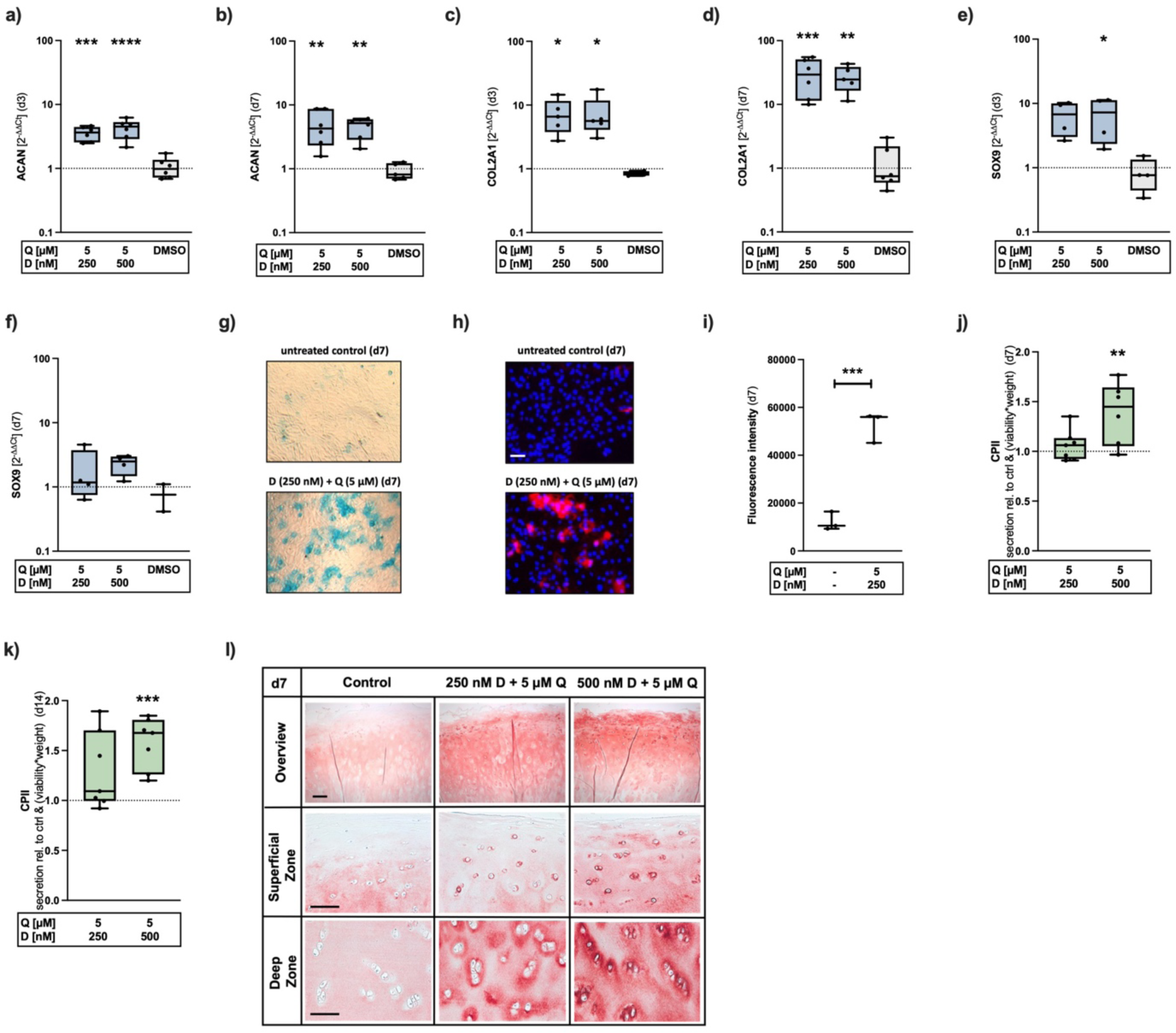
Promotion of the chondroanabolism after elimination of senescent hAC using D+Q. a+b) Gene expression levels of ACAN, c+d) COL2A1 and e+f) SOX9 during application (d3) and after withdrawal (d7) of the senolytics. g) Alcian blue staining of isolated hAC (d7), h) exemplary images of COL2-IF staining (scale bar = 100 µm) and i) the corresponding quantification of the FI. j+k) Relative CPII levels after senolytic therapy (d7+d14) and exemplary COL2 stainings (d7) of untreated human cartilage and after treatment with D+Q (scale bar: overview = 200 µm, superficial zone = 200 µm, deep zone = 100 µm). Blue colored box plots = gene expression analysis data; green colored box plots = data of protein analysis. Statistical analysis: a-e) one-way ANOVA with *Šidák*’s multiple comparisons test: *p<0.05, **p<0.01, ***p<0.001, ****p<0.0001; i) unpaired t-test ***p<0.001; j+k) one-way ANOVA with *Dunnett*’s multiple comparisons test: **p<0.01, ***p<0.001.

### D+Q Treatment Promotes *in vitro* Chondrogenesis of OA hAC but not ATDC5 Cells

As mentioned above, chondrocytes undergo phenotypical alteration through *in vitro* expansion, which is reflected in the loss of cartilage-typical extracellular matrix production. Therefore, we used high passage hAC, which can be considered as “dedifferentiated”, and studied their “redifferentiation” potential after D+Q treatment by means of an *in vitro* chondrogenesis assay. For this purpose, we treated hAC with D+Q for 3 d and compared the differentiation potential with the untreated hAC in a 3D pellet culture for 28 d. Based on the safranin-O and COL2 staining, we observed that elimination of senescent cells using D+Q promoted the chondrogenic differentiation capacities of the surviving hAC as demonstrated by a significant increase in the overall score (Fig. 4a+b). Next, we investigated, whether the treatment with D+Q can induce chondrogenic differentiation in ATDC5 cells. The gene expression analysis of COL2A1 (Fig. 4c) and ACAN (Fig. 4d) demonstrates that both were induced by CDM, while no increase of the chondroanabolic markers was observed after D+Q treatment in basal medium. These results were in line with the alcian blue and COL2-IF staining (Fig. 4e+f). Since chondrogenic differentiation in ATDC5 cells is known to be induced by ITS, D+Q treatment was performed in absence of ITS. To exclude that the presence of ITS was required in D+Q-related chondroanabolic response of hAC observed above, we cultured hAC in analogue conditions. In hAC, ITS administration displayed no beneficial effects on the gene expression levels of COL2A1 (Fig. 4g) and ACAN (Fig. 4h) or GAG production (Fig. 4i) as compared to ITS-free conditions, which suggests that the enhancement of the chondroanabolism in hAC can be attributed to D+Q treatment alone. Since the senolytic treatment promoted the chondrogenic phenotype in hAC but not in ATDC5 cells, we concluded that the therapy can be seen as a kind of reprogramming or restoring of the former phenotype rather than a direct inducer of chondrogenesis.

**Figure 4:**
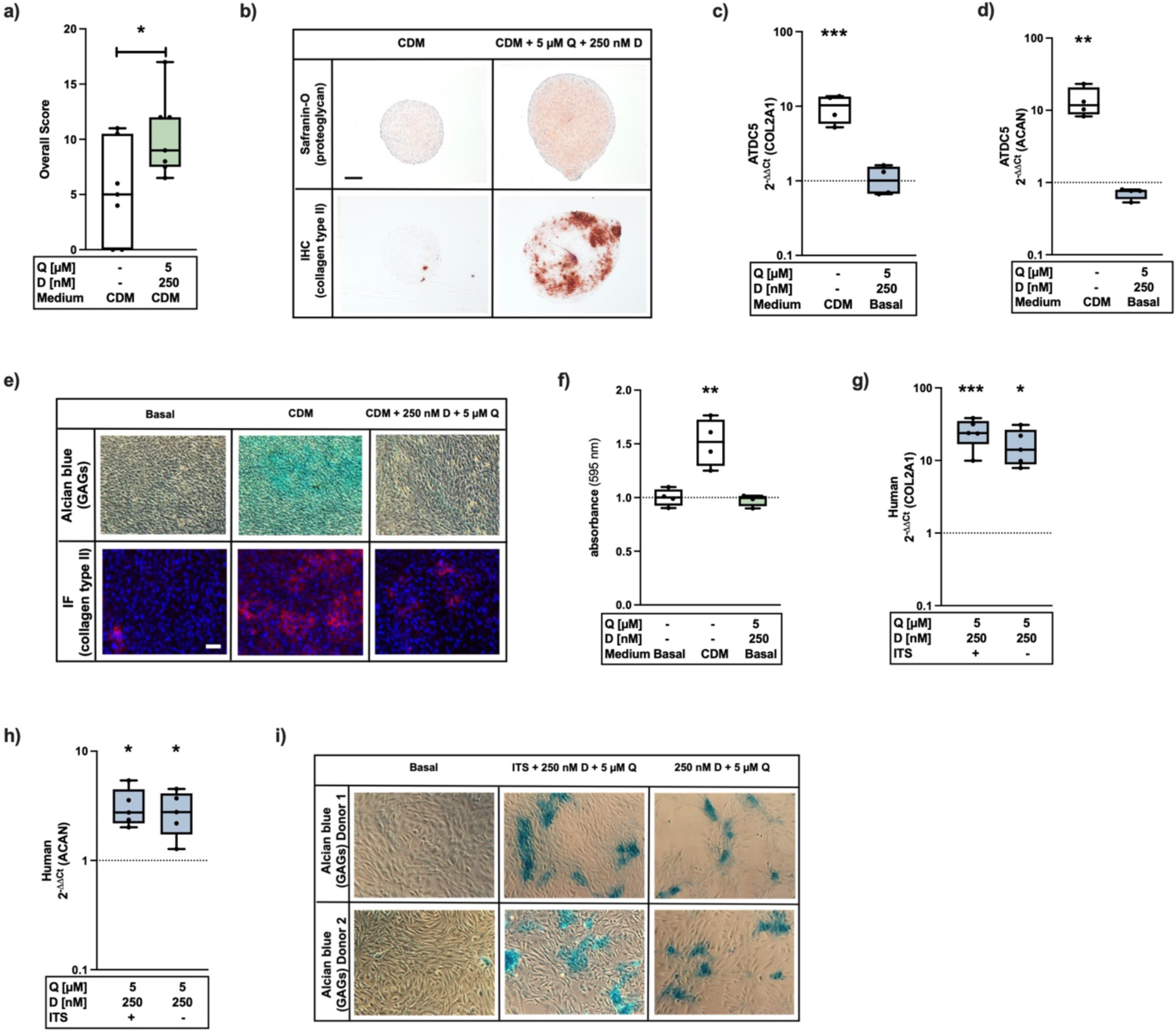
Influences of D+Q treatment on the chondrogenic phenotype of hAC and ATDC5 cells. a) Overall score of the chondrogenic redifferentiation potential of dedifferentiated hAC and b) exemplary images of the COL2 and safranin-O staining (scale bar = 200 µm). c) Gene expression analysis of COL2A1 and d) ACAN in ATDC5 cells after treatment with CDM or basal medium with D+Q. e) Corresponding alcian blue and COL2-IF staining (scale bar = 100 µm) of ATDC5 cells. f) Absorbance at 595 nm after destaining alcian blue with 1% SDS. g) Relative mRNA levels of COL2A1 and h) ACAN after senolytic treatment of hAC with or w/o ITS. g) Exemplary alcian blue staining of untreated hAC (basal) and after D+Q treatment with and w/o ITS. Blue colored box plots = gene expression analysis data; green colored box plots = data of protein analysis. Statistical analysis: a) unpaired t-test: *p<0.05; c+d) + f-h) one-way ANOVA with *Dunnett’s* multiple comparisons test: *p<0.05, **p<0.01, ***p<0.001.

### Single Treatment of Dasatinib but not Quercetin or Navitoclax Promotes Chondroanabolism in High Passage hAC

Next, we examined if both senolytics (D+Q) are required to achieve the promotion of the chondroanabolism or whether this can be achieved by single therapy of either D or Q. Based on the results obtained after separate application of either D or Q, Dasatinib was found to induce chondroanabolic effects, as shown by a strong increase of SOX9 (d3) (Fig. 5a) and COL2A1 (d7) (Fig. 5b) gene expression, as well as by an elevated production of GAGs verified by means of an alcian blue staining (Fig 5c). Quercetin did not affect the expression of chondroanabolic markers. One possible underlying mechanism of Dasatinib-mediated promotion of the chondrogenic phenotype could be through the inhibition of the Yes-associated protein (YAP-1) [44]. In our experiments, YAP1 mRNA levels were slightly downregulated by D or Q and the combination (Fig. 5d). As a next step, we clarified whether the clearance of senescent cells per se promotes the chondrogenic phenotype of high passage hAC or if this is related to the unique effects of Dasatinib. For this purpose, senescent cells were eliminated using Navitoclax, a commonly used and well-described senolytic, which induces apoptosis by targeting anti-apoptotic proteins of the BCL-2 family [32–34]. Navitoclax revealed stronger cytotoxic effects in high-passage (P≥4) hAC compared to low passages (P≤2) (Fig. 5e) (S4) and reduced the number of p16-positive cells in cartilage tissue (Fig. 5f) as similarly demonstrated for D+Q therapy above (Fig. 1d and Fig. 2h+i). However, in contrast to D+Q, gene expression levels of COL2A1 and ACAN of isolated high passage hAC were not elevated upon Navitoclax treatment (Fig. 5g-j). In addition, Navitoclax could not promote the chondrogenic redifferentiation potential of “dedifferentiated” high passage hAC (Fig. k+l). Overall, the results clearly demonstrate that the elimination of senescent cells alone does not promote chondroanabolism and that this effect is particularly related to Dasatinib.

**Figure 5:**
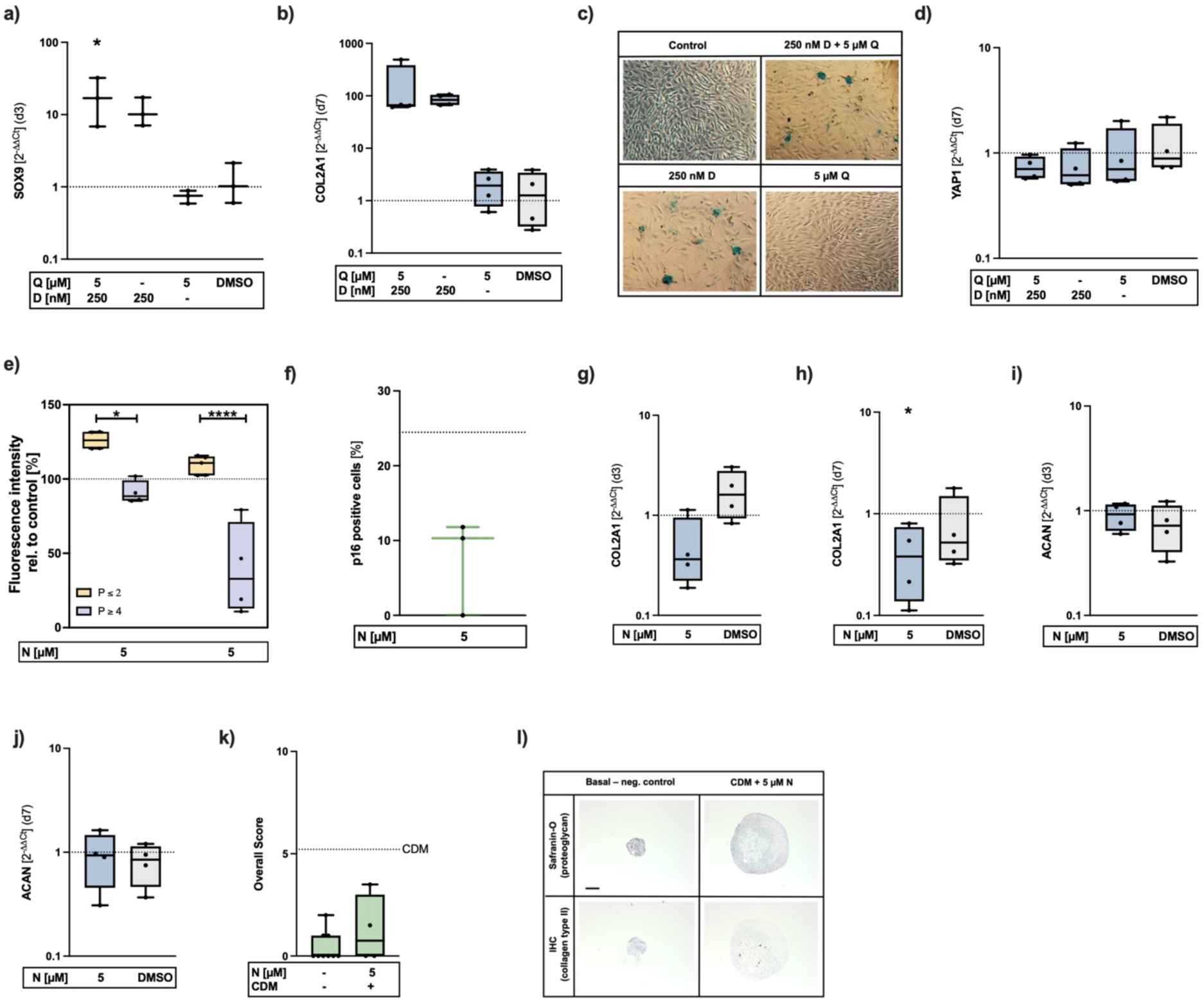
Dasatinib promotes chondroanabolism in hAC, while Navitoclax has impairing effects. a) mRNA levels of of SOX9 (d3) and b) of COL2A1 (d7) after stimulation with D, Q or D+Q. c) Exemplary alcian blue staining after single and combination therapy with D+Q. d) Gene expression levels of YAP1 after D, Q or D+Q therapy (d7). e) FI (alamarBlue assay) of low (P≤2) vs. high passage (P≥4) hAC after treatment with Navitoclax. f) p16 positive cells in cartilage tissue (IHC staining). g+h) Gene expression levels of COL2A1 and i+j) ACAN at d3+7 after Navitoclax therapy. k) Overall score of the chondrogenic redifferentiation potential after Navitoclax therapy of high passage hAC and l) exemplary images of the safranin-O and COL2 staining (scale bar = 200 µm). Blue colored box plots = gene expression analysis data; green colored box plots = data of protein analysis. Statistical analysis: a) one-way ANOVA with *Šidák*’s multiple comparisons test: *p<0.05; e) two-way ANOVA with *Šidák*’s multiple comparisons test: *p<0.05, ****p<0.0001; h) unpaired t-test: *p<0.05

### Senolytic Treatment Results in Downregulation of SASP Factors and Creates a Pro-Chondrogenic Environment

Senescent cells are characterized by a cell type-specific SASP. In case of chondrocytes, SASP factors contribute to a pro-inflammatory and anti-anabolic environment, which provokes a shift in cartilage homeostasis towards catabolism, while suppressing extracellular matrix synthesis. Furthermore, these mediators are assumed to promote increasing senescence in neighboring cells [1, 13, 16].

By the following experiment, we tested if the supernatant of senescent cells, containing various SASP factors, was able to affect the phenotype of low passage hAC with regard to senescent and chondroanabolic markers. For this purpose, we induced senescence in hAC by Doxo as previously described [12]. The senescent phenotype of Doxo-stimulated cells was confirmed on mRNA level (Fig. 6a+b). The SASP-conditioned medium of these cells was then used to stimulate non-senescent hAC for 48 h. Exposition to SIPS-conditioned medium reduced proliferation (Fig. 6c) and increased gene expression of CDKN1A (Fig. 6d), while suppression that of COL2A1 (Fig. 6g) in hAC as compared to hAC cultivated in non-conditioned medium. No differences were detected in case of CDKN2A and ACAN mRNA levels (Fig. 6 e+f). Similar results on COL2A1 gene expression were also achieved by stimulating cells with medium from high passage hAC (SASP-CM). Conversely, treatment of hAC with medium derived from cells prior stimulated with D+Q (DQ-CM) significantly increased the gene expression of COL2A1 (Fig. 6i) This indicated that Dasatinib promotes chondroanabolism via creation of a pro-chondrogenic environment. To confirm our suggestion, the expression of SASP markers was investigated in isolated high passage hAC directly after therapy (d3) and after deprivation (d7) of D+Q. The senolytic therapy resulted in a downregulation of the CXCL1 and IL6 gene expression (S5a+c), however, this effect was not maintained after deprivation of the senolytics (S5b+d). Similar results could be confirmed on protein level by means of an ELISA (Fig. 6j-m). The senolytic treatment had no influence on the release of IL8 (Fig. 6n+o). Since senescent chondrocytes secrete considerable amounts of MMPs, which in turn promote degradation of ECM components [14], MMP13 expression was examined. The results demonstrate that the mRNA levels were significantly reduced during therapy, which could be sustained even after withdrawal (S5e+f).

**Figure 6:**
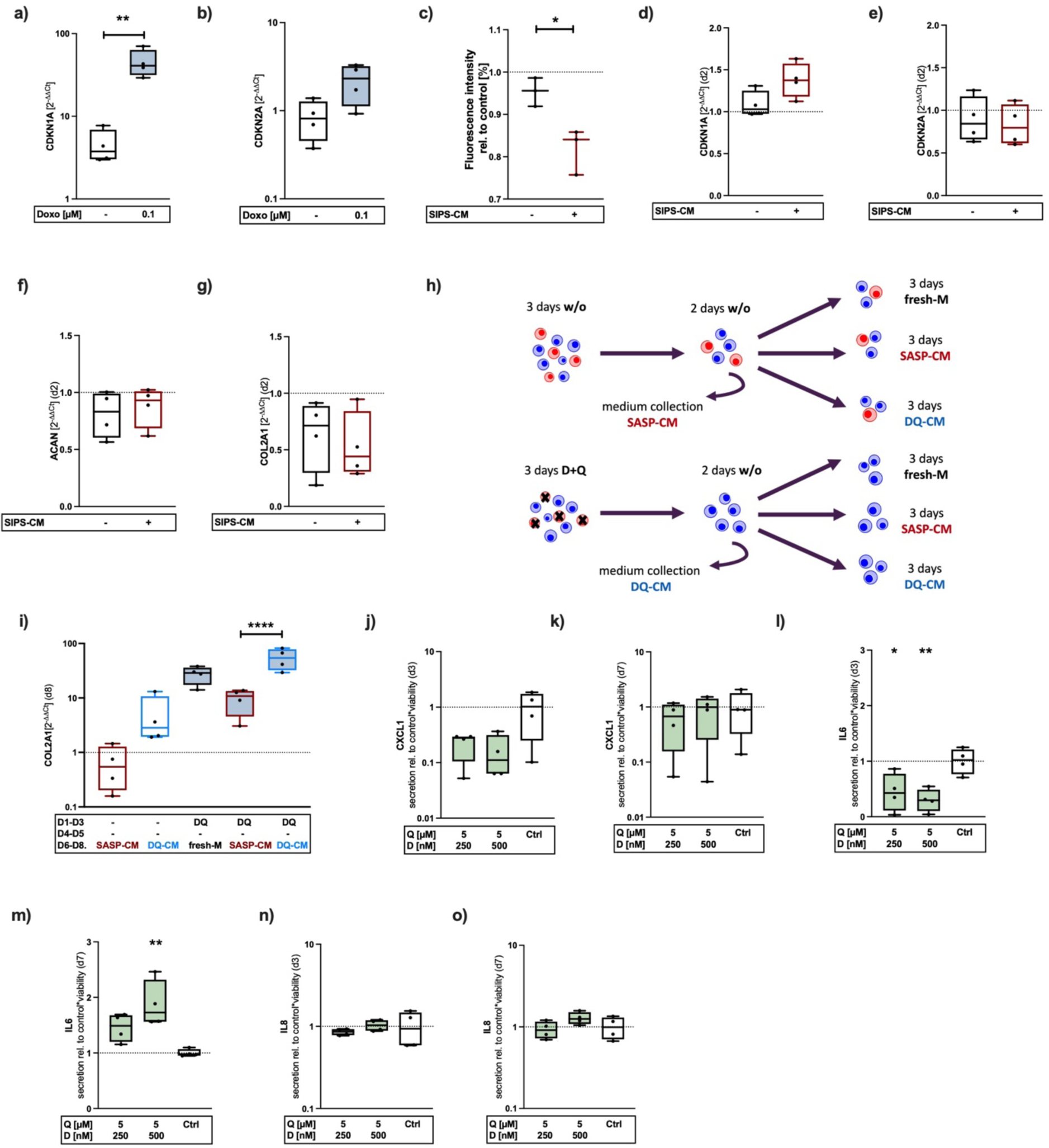
Analysis of hACs’ characteristic SASP factors and paracrine effects after D+Q treatment. a+b) mRNA levels of CDKN1A and -2A after stimulation with Doxo. c) alamarBlue assay: relative FIs after stimulation of hAC with SIPS-CM. Gene expression analysis of d) CDKN1A, e) -2A, f) ACAN and g) COL2A1 after stimulation with SIPS-CM. h) Experimental setup and i) relative mRNA levels of COL2A1 after stimulation of hACs with DQ-CM or SASP-CM. j+k) Quantification of secreted CXCL1, l+m) IL6 and n-o) IL8 at d3 and 7 by means of an ELISA. Blue colored box plots = gene expression analysis data; green colored box plots = data of protein analysis. Statistical analysis: a)+c) unpaired t-test: *p<0.05, **p<0.01; i)+l)+m) one-way ANOVA with *Šidák*’s multiple comparisons test: *p<0.05, **p<0.01, ****p<0.0001.

### RNA-Sequencing Reveals Downregulation of SASP Factors and Upregulation of Chondroanbolic Markers and Growth Factors

Total RNA-Sequencing analysis demonstrated a significant upregulation of 348 genes and a significant downregulation of 426 genes (Fig. 7a+b). In detail, increased gene expression was measured for several markers related to hyaline cartilage (COL2A1, COL9A1/2/3, COL11A1, MATN4, SOX9, and SNORC) and chondroanabolic growth factors (FGF18, IGF1, and TGFB2) (Fig. 7a+b). The upregulation of FGF18 and IGF1 was further confirmed via qRT-PCR (Fig. 7c+d). At the same time, anti-anabolic IGF1 binding proteins, IGFBP1 and 3, were significantly reduced (Fig. 7a+b) and the expression of FGF receptors (FGFR)-2, -3, and -4 was upregulated (Fig. 7a+b). Subsequent qRT-PCR analysis revealed a decreased FGFR1/FGFR3 expression ratio through senolytic therapy (Fig. 7e), implying a shift towards chondroanabolic FGFR-signaling. In line with this, gene expression of SERPINA1 and SERPINA3, encoding for alpha1-antitrypsin (AAT), was significantly enhanced (Fig. 7a+b).

**Figure 7:**
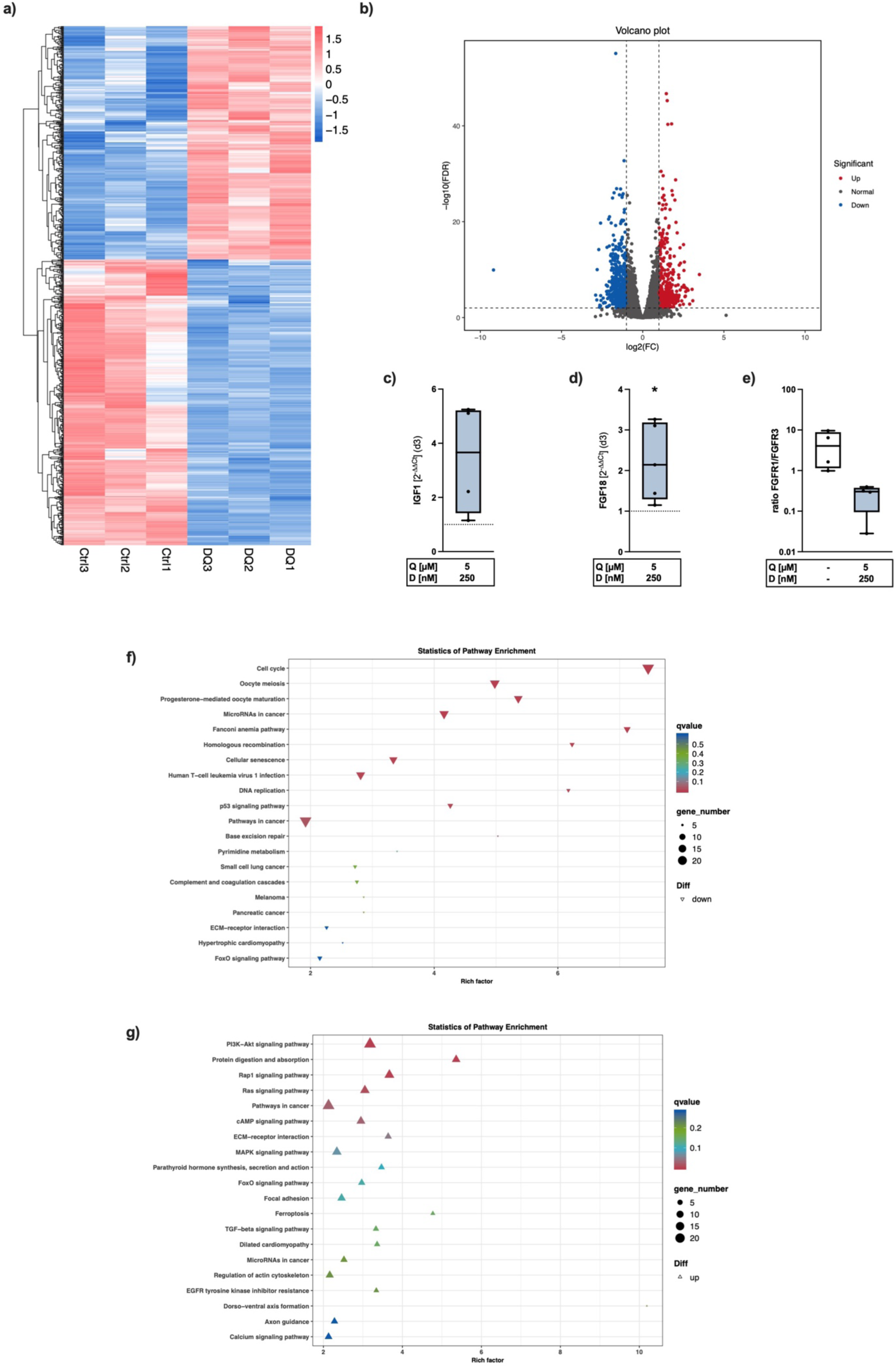
RNA-sequencing analysis after D+Q treatment. a) DEG-Plot – significantly up and downregulated genes. b) Volcano Plot – significantly up- and downregulated genes. Confirmation of RNA-seq data by qRT PCR analysis of c) IGF1, d) FGF18, and e) FGFR1/3 ratio (d3). Results of KEGG pathway enrichment analysis of f) down- and g) upregulated signaling pathways. Statistical analysis: a+b), f+g): t-test, d) unpaired t-test.

CTGF, BIRC5 (survivin), and CYR61, target genes of YAP-1 [45], were downregulated, thus indicating an inhibition of the YAP-1 signaling pathway (Fig. 7a+b). Additionally, the therapy resulted in an increased gene expression of the extracellular antioxidative enzyme SOD3 (Fig. 7a+b). Further anti-inflammatory properties were displayed by the decreased expression of IL6, IL7R, IL26, IL32 (Fig. 7a+b). Anti-catabolic effects were demonstrated by downregulation of the aggrecanases ADAMTS-1 and 5 (Fig. 7a+b). Furthermore, the treatment decreased the expression level of RIPK3 (Fig. 7a+b), an important mediator during necroptosis [46]. In addition, the senolytic therapy resulted in the downregulation of several genes related to proliferation (Fig. 7a+b).

Overall, the Kyoto Encyclopedia of Genes and Genomes (KEGG) pathway enrichment analysis revealed a downregulation of genes associated with cell cycle, cellular senescence, DNA replication, and p53 signaling (Fig. 7f). An upregulation was detected in case of the PI3K-Akt-, and TGF-β-signaling pathway (Fig. 7g). Moreover, genes related to the FoxO signaling pathway, which plays an important role in chondrocyte function and senescence [47], were significantly regulated upon D+Q treatment (Fig. 7f+g).

## Discussion

The relevance of research into new therapies for the treatment of OA becomes particularly clear, given that 654.1 million people worldwide are affected by osteoarthritis of the knee joint, as estimated in 2020 [48]. Currently, no disease modifying OA drugs (DMOADs) are approved and therefore OA is still an uncurable disease [49, 50]. Due to the detrimental effects of cellular senescence on tissue homeostasis, selective elimination of senescent cells received increasing attention in OA as well as other degenerative diseases during the last decades [32, 51]. Although the senolytic UBX0101 failed in phase II clinical trials as a potential therapeutic agent in knee OA [32, 51], senolysis is still considered a promising strategy and further senolytics, like fisetin, are currently under investigation in OA patients (NCT04210986).

In our study, we tested the senolytic combination therapy with D+Q for the first time in human cartilage tissue derived from OA patients. Considering our novel findings on D+Q therapy, we concluded that the senolytics could be a suitable therapeutic approach against OA in many ways. D+Q selectively eliminated senescent chondrocytes in cell as well as tissue culture. The treatment resulted in a tremendous promotion of the chondroanabolism in surviving chondrocytes, demonstrated by elevated COL2 and PG levels. Additionally, D+Q treatment of hACs improved the chondrogenic redifferentiation indicating a rejuvenation or cellular reprogramming of the dysfunctional cells. Overall, our results imply that D+Q therapy, and in particularly Dasatinib, restored the chondrogenic phenotype of hAC, while elimination of senescent cells by Navitoclax had no beneficial effect regarding the chondroanabolism. Furthermore, we provide evidence that D+Q-mediated senolysis created a pro-chondrogenic environment by suppressing the production of pro-inflammatory and anti-anabolic SASP factors, such as CXCL1, IL6, and IGFBPs, and increasing the expression of chondroanabolic growths factors.

After identifying the beneficial effects of D+Q treatment on the chondroanabolism, we investigated if D+Q in combination is required or if this can be achieved by single therapy of one of those. In detailed analyses, we demonstrated that Dasatinib alone promoted the chondrogenic phenotype. Several *in vivo* studies provide evidence that quercetin may be useful for the treatment of OA, particularly in the prevention of PTOA [52–54]. Administration of quercetin has been described with anti-nociceptive [54], cell- and chondroprotective [53, 55] as well as anti-inflammatory properties [55], thus attenuating surgically-induced OA progression in rats. Monotherapeutic treatment with Dasatinib, in turn, exhibited pro-regenerative effects in a rabbit full-thickness cartilage defect-model [44]. In fact, it has been reported that Dasatinib promotes MSCs chondrogenesis, presumably via the inhibition of the Src/Hippo-YAP signaling pathway. The treatment resulted in a decreased expression and enhanced cytoplasmic degradation of YAP [44]. The downregulation of YAP is known to be necessary for the chondrogenesis of MSCs [44, 56], while high YAP activity, in contrast, is thought to suppress chondrogenic differentiation [56]. Moreover, the YAP-TEAD complex was found to maintain survival of senescent cells [57]. In our study, we could confirm a significant decrease of the three main signature genes of the YAP1-signaling pathway (BIRC5, CTGF, and CYR61) upon D+Q treatment by means of an RNA-sequencing. Furthermore, we observed a negative trend in the transcription of YAP1 itself. Although, Quercetin had no noticeable beneficial effect regarding the pro-anabolic properties of Dasatinib, we could not find any adverse effect either. Considering this and the beforementioned positive effects of Quercetin reported in other studies [52–55], we suggest that the combination of D+Q is still reasonable. Next, we considered whether the elimination of senescent cells alone exerts beneficial effects on the chondroanabolism. Therefore, we treated chondrocytes with Navitoclax and confirmed the selective elimination of senescent cells through this therapy. However, Navitoclax could not support the chondroanabolism but rather impaired the chondrogenic phenotype of hAC. In contrast, Yang et al. reported not only chondroprotective effects in a rat PTOA model, but also an increase in the chondrogenic phenotype of human OA chondroyctes after Navitoclax treatment [34]. Nevertheless, there are further studies from the musculoskeletal field, which observed anti-anabolic effects of Navitoclax, confirming our findings. For example, Sharma et al. reported trabecular bone loss and impaired osteoprogenitor cell function in osteoporotic mice after Navitoclax treatment [58]. In contrast, D+Q therapy had anabolic and anti-resorptive effects on bone, thus preventing bone loss and enhancing the repair potential in murine postmenopausal and age-related osteoporosis models [20–22]. Although D+Q therapy displayed chondroprotective and chondroanabolic properties in OA as described above [19], the combination of D+Q with Navitoclax had, interestingly enough, no modifying effect on cartilage degeneration in a model of spontaneous OA [59]. Overall, these findings imply that senolysis alone does not restore the phenotype of bone or cartilage cells and that senolytics have to be carefully chosen, depending on the target tissue and cell type.

Further, we were able to demonstrate that cells which received medium from previously D+Q stimulated cells expressed higher levels of COL2A1, while medium from untreated high passage cells had suppressing effects. We supposed that this could be due to the creation of a pro-chondrogenic microenvironment through D+Q therapy. In line with our assumption, the results of the RNA sequencing analysis provided evidence that the chondroanabolic effects of the conditioned medium might be explained by upregulation of relevant growth factors (IGF1, FGF18, TGFB2) as well as SERPINA1/3 (ATT) and concomitant reduction of anti-anabolic proteins, including cytokines and IGFBPs. IGF1 and FGF18 are well-known drivers of the ECM synthesis [60]. In a previous study, we could confirm the chondroanabolic effects of IGF-1 on cartilage tissue by an increased COL2A1 gene expression levels and elevated CPII levels, a biomarker for COL2 synthesis [61]. FGF-18 treatment displayed pro-anabolic features in various animal studies. For example, it enhanced cartilage repair in an ovine model after a chondral defect [62] and displayed cartilage protective effects in a rat PTOA model [63].

Additionally, we previously observed cell- and chondroprotective effects of FGF18 therapy [61]. In case of ATT, it has been reported that SERPINA1 and 3 are highly expressed in articular cartilage and during chondrogenesis of MSCs, but decreased during *in vitro* expansion (dedifferentiation) of hAC and OA progression [64]. Furthermore, AAT exerted chondroprotective and chondroanabolic effects [65]. Altogether, increased expression of markers associated with hyaline cartilage formation (e.g., ACAN, COL2A1, COL9A1/2/3, COL11A1, MATN4), chondrocyte maturation (e.g., SNORC, SOX9, SERPINA1/3), and the observed decline in proliferation in D+Q-treated hAC, imply that the dedifferentiated hAC return to a mature, non-mitotic state. This is very impressive regarding the previous assumption that serum-free conditions and a 3D environment are obligatory requirements in redifferentiation of primary chondrocytes [66, 67]. Beyond the pro-anabolic effects, the sequencing analysis revealed a downregulation of aggrecanases and pro-inflammatory factors after D+Q treatment. Altogether, the reduction of SASP factors is of great importance, as they negatively influence the surrounding cells [68]. However, it should be noted that our data imply that senolysis per se does not ensure a pro-chondroanabolic microenvironment, as Navitoclax rather suppressed COL2 synthesis despite elimination of senescent.

The therapy with D+Q is currently investigated in clinical trials regarding its diseases modifying effects in common age-related disorders such as the chronic kidney disease or the idiopathic pulmonary disease [32, 69–71]. Therapy with D+Q was generally well-tolerated [69–71], which suggests that intraarticular injection of D+Q in OA joints would be similarly innocuous and could – with regard to our meaningful *ex vivo* results – certainly be a promising therapeutic approach in patients suffering from knee OA. Despite previous *in vitro* and *in vivo* investigations on the effects of D+Q in OA, we consider our study as novel and relevant. However, previous studies did not investigate the phenotype of surviving chondrocytes. Moreover, the results derived from animal experiments, mainly performed on young animals, cannot be directly transferred to the clinical situation in which predominately elderly people suffering from OA. Therefore, we used human OA cartilage, that provides important information about the efficacy of D+Q on aged and OA-affected tissue. This tissue contains senescent cells which have accumulated in the course of OA progression and were not provoked by extrinsic manipulation, such as radiation [72] or addition of chemicals [12], thus perfectly mirroring the clinical appearance. Despite of the advantages of using patient material, our study is limited to the effects on cartilage tissue and does not consider other joint-related tissues, such as the synovial membrane or subchondral bone, involved in OA progression. Moreover, it was not possible to include a healthy control group in our study. To overcome this issue, healthy porcine tissue had to be used.

Considering our results, we could demonstrate that promotion of the chondrogenic phenotype by D+Q after selective elimination of senescent chondrocytes might be based on two mechanisms: first, direct modulation of the YAP-1 signaling pathway and, second, paracrine effects through reduction of detrimental SASP factors and increased expression of pro-chondroanabolic mediators. Altogether, this senolytic combination therapy could offer a new therapeutic strategy in OA.

## Author Contribution

SM: conceptualization, experiment conduction and data analysis, visualization, manuscript writing. JRK: funding acquisition, conceptualization, experiment conduction and data analysis, visualization, manuscript writing. VK+LR: experiment conduction and data analysis, review and editing of manuscript. RB: review and editing of manuscript.

## Acknowledgements

Special acknowledgements are given to Natalie Braun, Christiane Schulz and Giovanni Ravalli for their technical assistance, and Jana Hengge for her laboratory work. This research study was supported by the European Social Fund and by the Ministry of Science, Research and Arts Baden-Württemberg, as well as the DFG (CRC1149 start-up funding and gender equality funding).

