## Supplementary material for "Senolytic therapy combining Dasatinib and Quercetin restores the chondrogenic phenotype of human osteoarthritic chondrocytes by promoting the release of pro-anabolic mediators": S

**Materials and Methods:**

**S1: Materials and Methods**

**a) Medium:**

- Serum-free medium – human:
  DMEM 1 g/L glucose (Gibco Life Technologies), 1% Na-pyruvate (Merck Biochrom), 1% non-essential amino acids (NEAAs) (Merck Biochrom), 2 mM L-glutamine (PAN-Biotech), 0.5% penicillin/streptomycin (PAN-Biotech), 10 µg/mL 2-phospho-L-ascorbic acid trisodium salt (Sigma-Aldrich), 1% 100x insulin-transferrin-selenium (ITS) (Gibco, Life Technologies)
- Serum-free medium – porcine:
  DMEM 4.5 g/L glucose (Gibco Life Technologies), 1% Na-pyruvate, 1% NEAAs, 2 mM L-glutamine, 1% penicillin/streptomycin, 10 µg/mL 2-phospho-L-ascorbic acid trisodium salt, 1% 100x ITS
- Serum-containing medium – human:
  1:1 Ham's F12 (PAN-Biotech) and DMEM 1 g/L glucose, 10% heat inactivated fetal bovine serum (FBS) (PAN-Biotech), 0.5% penicillin/streptomycin, 2 mM L-glutamine, 10 µg/mL 2-phospho-L-ascorbic acid trisodium salt
- Chondrogenic differentiation medium – human:
  DMEM 4.5 g/L glucose, 1% penicillin/streptomycin, 2 mM L-Glutamine, 0.1 µM Dexamethasone (Sigma-Aldrich), 0.2 mM 2-phospho-L-ascorbic acid trisodium salt, 40 µg/mL L-proline, 1 mM Na-pyruvate, 10 ng/mL of rhTGF-$\beta$3 (PeproTech) and 10 ng/mL rhBMP-6 (PeproTech), 1% 100x insulin-transferrin-selenium (ITS) (Gibco, Life Technologies)
- Serum-containing medium – ATDC5 cells:
  1:1 DMEM 1 g/L glucose, HAM's F12, 5% FBS, 1% L-glutamine, 10 µg/mL Transferrin human (Sigma Aldrich), 0.03 µM Sodium Selenite (Sigma Aldrich), 1% penicillin/streptomycin
- Chondrogenic differentiation medium – ATDC5 cells:
  DMEM 4.5 g/L glucose, 1% penicillin/streptomycin, 5% FBS, 2 mM L-Glutamine, 0.1 µM Dexamethasone (Sigma-Aldrich), 0.2 mM 2-phospho-L-ascorbic acid trisodium salt, 40 µg/mL L-proline, 1 mM Na-pyruvate, 10 ng/mL of rhTGF-$\beta$3 (PeproTech) and 10 ng/mL rhBMP-6 (PeproTech), 1% 100x insulin-transferrin-selenium (ITS) (Gibco, Life Technologies)

**b) Probes and Primers:**

- Probes – human:
  Reference genes: GAPDH (Hs02758991_g1), HPRT1 (Hs02800695_m1), and TMEM199 (Hs01022209_m1)
  Target genes:
  COL2A1 (Hs00264051_m1), ACAN (Hs00153936_m1), CXCL1 (Hs00605382_gH), IL6 (Hs00174131_m1), MMP13 (Hs00233992), CDKN1A (Hs00355782_m1), CDKN2A (Hs00923894_m1), YAP1 (Hs00902712_g1) SOX9 (Hs00165814_m1), FGFR1 (Hs00241111_m1), FGFR3 (Hs00179829_m1)
- Probes and primers – human:

Target genes: IGF1 (forward 5'CTCTTCAGTTCGTGTGTGGAGAC; reverse 5'CAGCCTCCTTAGATCACAGCTC3'), FGF18 (all in one BioCat HQP088256-GC)

- Probes and primers – murine:
  Reference genes: GAPDH (Mm99999915_g1), HPRT1 (Mm03024075_m1)
  Target genes: ACAN (forward 5'AACTTCTTTGCCACCGGAGA3'; reverse 3'GGTGCCCTTTTTACACGTGAA5'), COL2A1 (forward 5'CCTGTCTGCTTCTTGTAAAAC3'; reverse 3'TGGGTATCATCAGGTCAGGT5').

**S2: Morphological Changes of hAC through Passaging**

Morphological changes due to *in vitro* culturing are demonstrated by a correlation of cell length with increasing passage.

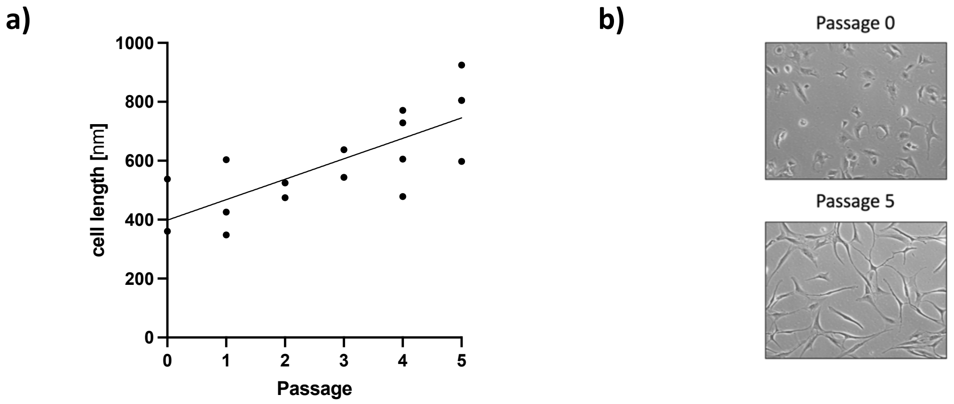


S2: a) Passage dependent cell length (passage 0-5) and b) exemplary images of passage 0+5.

**S3: Enhanced Chondroanabolism in Cartilage Tissue through D+Q Therapy**

COL2 staining intensity of cartilage tissue was increased after senolytic treatment with D+Q at day 14.

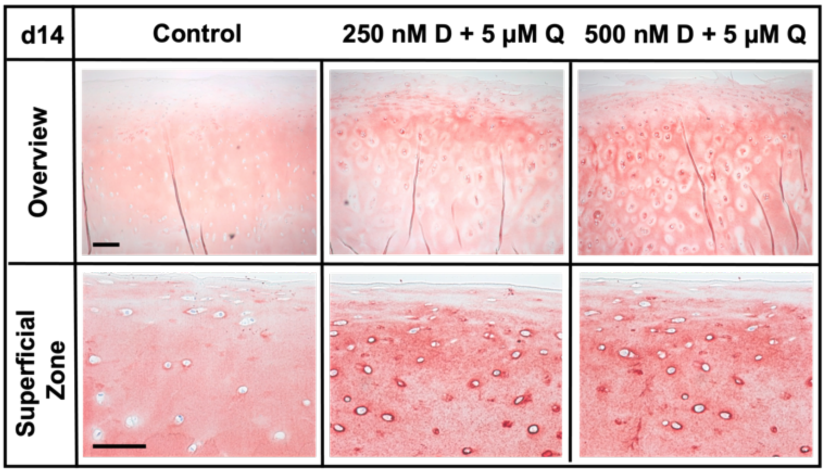


S3: Exemplary COL2 staining (d14) of cartilage tissue w/o or with D+Q therapy (scale bar: overview = 200 µm, superficial zone = 200 µm).

**S4: Cytotoxic Effects of Navitoclax on Isolated hAC**

Caspase 3/7 activity relative to the untreated control in low. vs. high passages as well as in healthy (OARSI ≤1) and degenerated (OARSI ≥3) cartilage tissue after senolytic therapy with Navitoclax (2.5-5 µM).

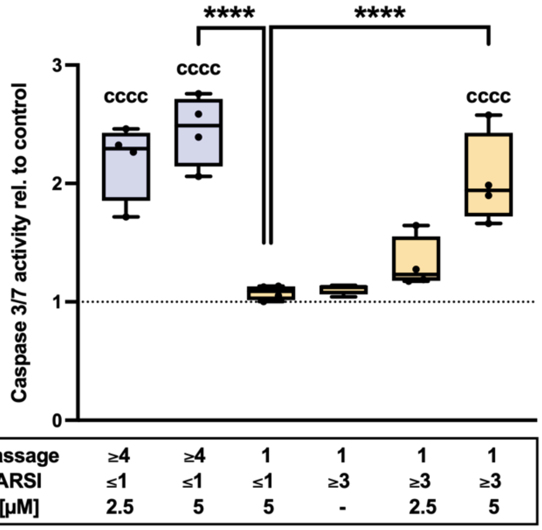


S4: Caspase 3/7 activity of high vs. low passage hAC w/o or with Navi (2.5-5 mM) treatment. Statistical analysis: one-way ANOVA with Šidák*'s* multiple comparisons test: ****p<0.0001.

**S5: Downregulation of SASP Factors on Gene Expression Level after Therapy with D+Q**

Senolytic treatment resulted in a downregulation of the gene expression levels CXCL1 and IL6 directly after therapy (d3). After deprivation of D+Q the mRNA levels were no longer downregulated. The protease MMP13 gene expression level was downregulated at d3+7.

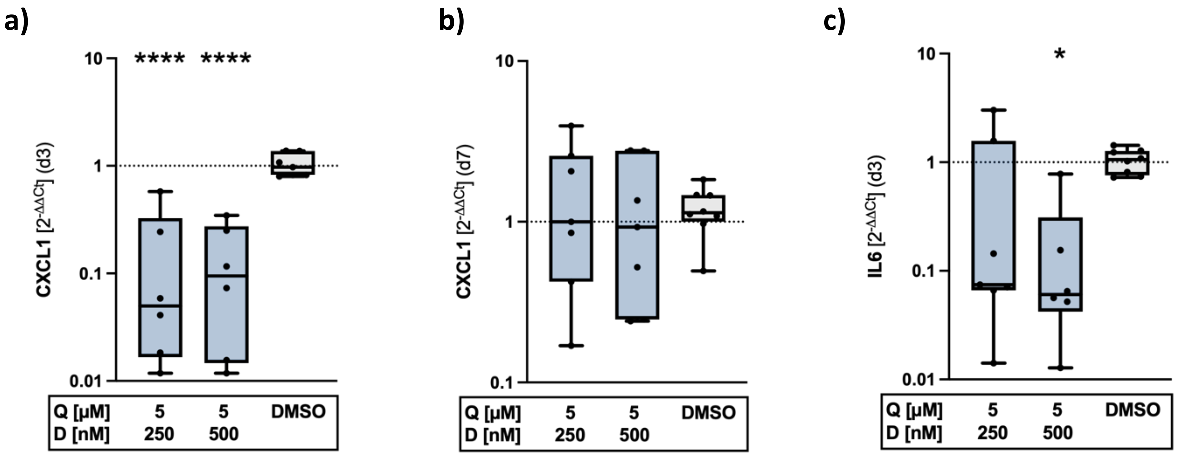

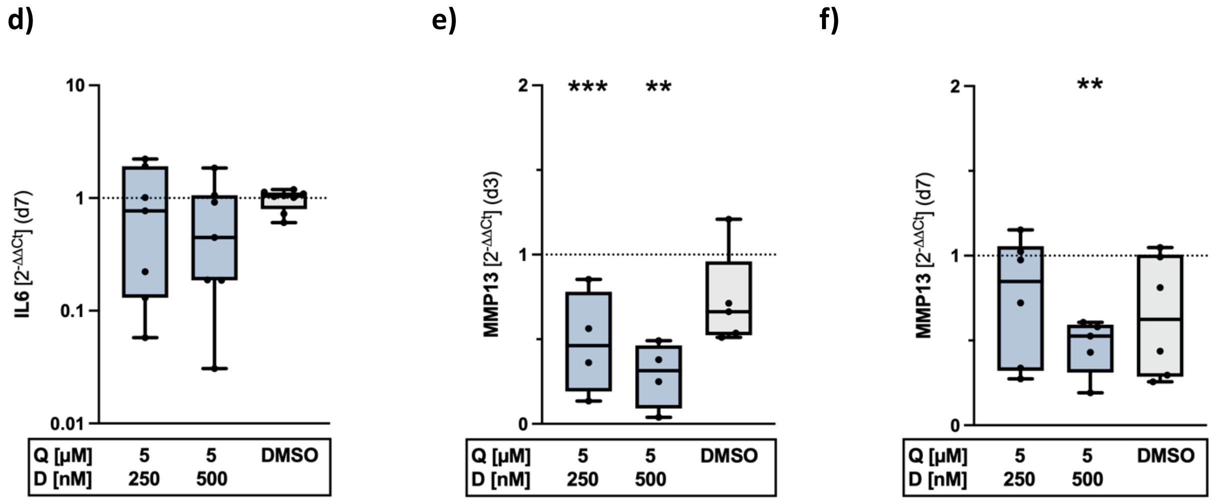


S5: Downregulation of SASP factor expression. a+b) Gene expression analysis of CXCL1 (d3+d7), c+d) IL6 (d3+d7), and e+f) MMP13 (d3+d7). Statistical analysis: one-way ANOVA with Šidák's multiple comparisons test: *p<0.05, **p<0.01, ***p<0.001, ****p<0.0001.
